# Activation of hypothalamic-pontine-spinal pathway promotes locomotor initiation and functional recovery after spinal cord injury in mice

**DOI:** 10.1101/2024.11.16.621796

**Authors:** Chengyue Ji, Yunfan Zhang, Zeyu Lin, Ziqi Zhao, Zhuolei Jiao, Zhiyuan Zheng, Xiaoxue Shi, Xiaofei Wang, Ziyu Li, Shuisheng Yu, Yun Qu, Yaxuan Wei, Bowen Zheng, Hanyu Shi, Qifang Wang, Alfredo Sandoval, Bo Chen, Xiao Yu, Xiaohong Xu, Mu-ming Poo, Juxiang Chen, Weihua Cai, Yi Li

## Abstract

The hypothalamus is critical for regulating behaviors essential for survival and locomotion, but how it integrates internal needs and transmits locomotion commands to the spinal cord (SC) remains unclear. We found that glutamatergic neurons in lateral hypothalamic area (LHA) are essential for regulating motivated locomotor activity. Using single-neuron projectome analysis, trans-synaptic tracing, and optogenetic manipulation, we showed that LHA facilitates motivated locomotion during food seeking via pontine oral part (PnO) projection neurons, rather than direct SC projections or indirect stress signaling via medial septum and diagonal band. Activating PnO-SC projection neurons also initiated locomotion. Importantly, LHA-PnO projection neurons were crucial for regulating locomotor recovery following mouse spinal cord injury (SCI). Closed-loop deep brain stimulation (DBS) of LHA via gating by motor cortex signals markedly promoted long-term restoration of hindlimb motor functions after SCI. Thus, we have identified a hypothalamic-pontine- spinal pathway and the stimulation paradigm for potential therapeutic intervention after SCI.

## INTRODUCTION

Locomotion is a fundamental behavior essential for survival, enabling animals to seek food, evade threats, and engage in social and reproductive activities. Two primary command centers in the brain, the subthalamic locomotor region (SLR) and the mesencephalic locomotor region (MLR), are responsible for transmitting locomotor commands to the central pattern generators in spinal cord ^1–4^. Extensively studies have revealed the role of MLR in controlling locomotion speed and various aspects of body movement ^5–8^, but the mechanisms by which the SLR integrates internal/external stimulation and transmits locomotor commands to the spinal cord remain unclear.

The lateral hypothalamus area (LHA) within the SLR has been implicated in motor control, particularly in motivated locomotion—internal need-driven movements such as food-seeking, exploration, or goal-directed behaviors ^9,10^. However, the precise LHA circuits controlling the locomotion, particularly within the contexts of motivation, remain incompletely understood ^3,11^. Recent single-neuron projectome studies have begun to reveal more detailed connections between specific neuronal subtypes in distinct brain regions, providing new insights into how these circuits may coordinate complex behaviors ^12–16^.

Defining the neural circuits that control locomotion is also crucial for developing neural circuit-based strategies to promote motor recovery following injuries of the central nervous system (CNS). This is particularly relevant in the case of SCI, in which the brain loses its direct connection with the SC. Despite neuronal death and axon degeneration after SCI, some axons and dormant relay pathways often remain spared across the lesion site, providing a neural substrate that can be reactivated to promote functional recovery, even with limited axon regeneration ^17^. The restoration of locomotor function after SCI involves the reorganization of both spinal and supraspinal circuits ^18^. Recent studies have unraveled brain region-specific contributions to motor function recovery after SCI, further arguing the importance of reactivation of the neural circuits involved in the regulation of locomotion control ^19–22^. However, effective therapeutic strategies for restoring motor function after SCI remain limited. As a promising strategy for in vivo neuronal activation, deep brain stimulation (DBS) offers the advantage of locally activating specific brain nuclei and has been successfully used to treat Parkinson’s disease ^23^. Recent studies have also revealed that targeting specific MLR regions, including cuneiform nucleus (CnF) ^24–26^ and pedunculopontine nucleus (PPN) ^27^, can enhance motor function and promote recovery in animal SCI models. However, the precise brain nuclei to target for optimal motor recovery remain under investigation ^28^, and clinical trials are still in their early stages ^29^.

Here, we employed a range of advanced techniques, including whole-brain trans-synaptic labeling, single-neuron projectome analysis, and projection-specific targeting, to investigate the contributions of the hypothalamic LHA nucleus in locomotion control, and in motor function restoration following SCI. Our findings suggest that LHA facilitates motivated locomotion through an indirect pathway, particularly the pontine reticulospinal tract, rather than through direct projections to the spinal cord or through connections to the medial septum and diagonal band (MSDB). This indirect modulation of spinal circuits offers a novel target for therapeutic intervention. Moreover, we investigated the potential of DBS targeting the LHA to promote motor recovery, with the ultimate goal of developing a translatable strategy for restoring locomotion in individuals with severe SCI.

## RESULTS

### LHA glutamatergic neurons are involved in hindlimb motor control

To visualize the hypothalamic areas that directly or indirectly projected to the hindlimb muscles, we first injected pseudorabies virus (PRV) encoding EGFP directly into the tibialis anterior/gastrocnemius (TA/GS) muscle. This allowed for retrograde trans-synaptic labeling of upstream neurons. The ClearMap ^30,31^ method was utilized for automatic analysis and registration of volumetric data from cleared tissues at 5 days after viral injection (see STAR Method). The reconstructed whole-brain data of labelled neurons were registered onto the standard Allen CCFv3 map, showing a widespread distribution of PRV-labeled neurons across the brain (Figure 1A). Further analysis showed that labeled neurons were widely distributed across the hypothalamus, with the highest density (36.6 % of the total) in the lateral hypothalamus area (LHA) (Figures 1B-D and S1A-B).

**Figure 1.**
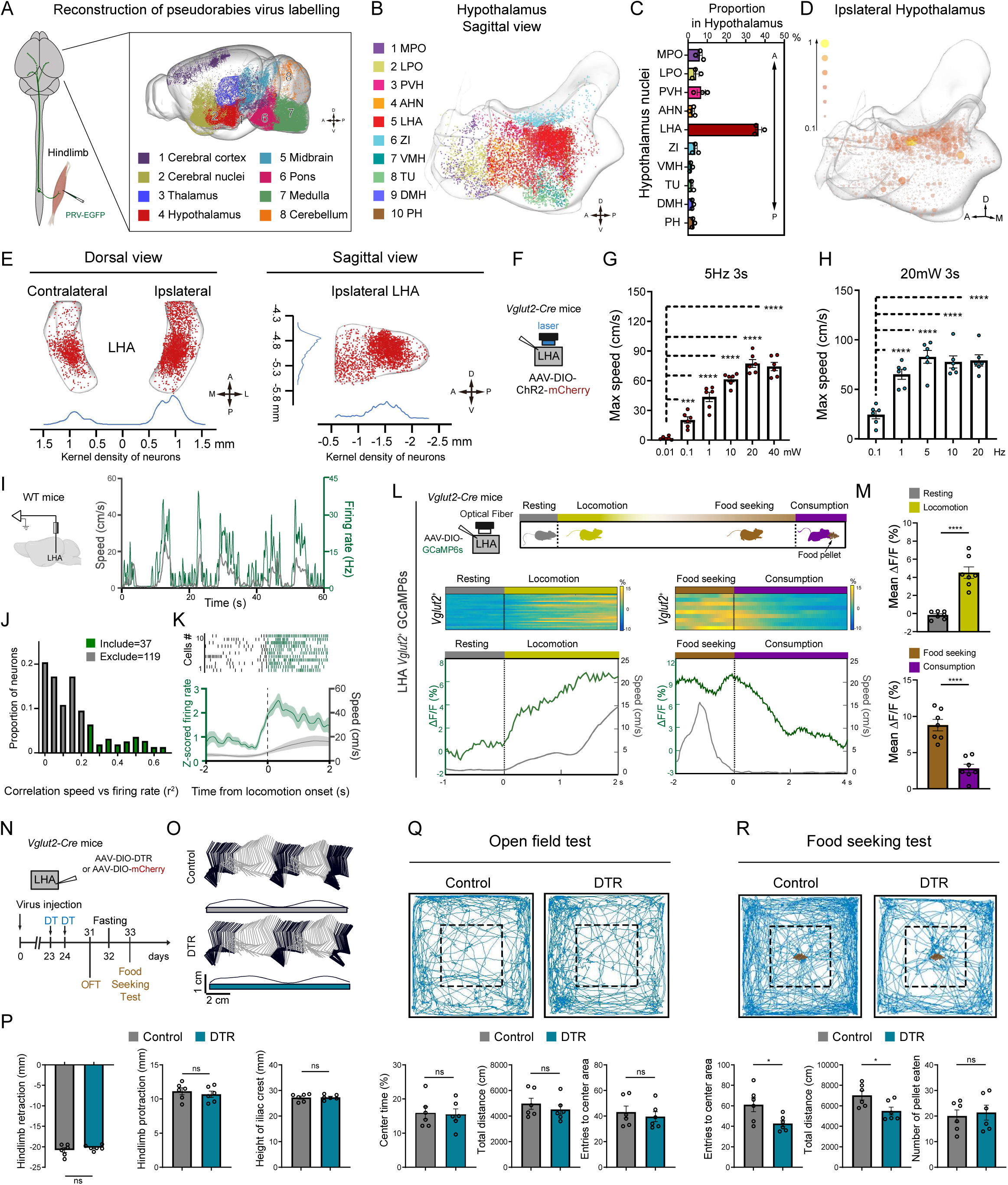
LHA Glutamatergic Neurons in the Hypothalamus Are Predominantly Involved in Hindlimb Motor Control and Required for Motivation-Driven Locomotion. (A) Left: Schematic of PRV-EGFP injection into the TA (tibialis anterior muscle) and GS (Gastrocnemius muscle). Right: Reconstruction of adult wild-type mouse brain labeled for EGFP after a viral delivery of PRV in the hindlimb muscle. (B) Reconstruction of PRV-EGFP labelled neurons in the hypothalamus innervating the hindlimb muscles. (C) Three-dimensional heatmap distribution of PRV-labeled cells within the hypothalamus. (D) The percentage of labelled neurons in the hypothalamus. (E) The dorsal view (left) and sagittal view (right) of PRV-infected lateral hypothalamus area neurons. (F) Strategy for activating LHA glutamatergic neurons. (G and H) Quantification of maximum speed corresponding to optical stimulation in different light intensities and frequencies. One-way ANOVA, followed by Bonferroni post hoc test. ***p < 0.001; ****p < 0.001. Error bars, SEM. (I) Left: Schematic of tetrode recording for LHA neurons and offline spike sorting. Right: Firing rate of one representative neuron in LHA (green line, right axis) plotted with the speed (gray line, left axis) of the mouse. (J) Distribution of the recorded LHA neurons showing the speed of locomotion based on their relative correlation (R^2^) with speed. Green bars, speed-correlated neurons; Gray bars, neurons show no significant correlation with the speed. (Spearman correlation test, P>0.05). (K) Top: Raster plot for spike responses of representative neurons to locomotion onset. Bottom: population z-scored firing rate of a subpopulation LHA neurons (green line) and speed (gray line) aligned to the onset of locomotion. (L) Top: Strategy for fiber photometry recording of LHA *Vglut2^+^* neurons and food seeking test in the runway. Bottom: Calcium dynamics of LHA *Vglut2^+^* neurons in response to onset of locomotion and consumption. (M) Quantification of LHA *Vglut2^+^* neurons calcium response to onset of locomotion (top) and food consumption (bottom). n = 6 mice, 3-5 trials per mouse. Student’s t test (two-tailed, unpaired) was applied. ****p < 0.0001. Error bars, SEM. (N) Timeline summarizing the experiments of gait analysis, open field and food seeking test with or without depletion of LHA glutamatergic neurons. (O) Representative stick diagram decomposition of leg movements in mice with or without depletion of LHA glutamatergic neurons. (P) Quantification of retraction, protraction and height of iliac crest with or without depletion of LHA glutamatergic neurons (n=6). Student’s t test (two-tailed, unpaired) was applied. ns, not significant. Error bars, SEM. (Q) Top: Trajectory of mice in open-field chambers with or without depletion of LHA glutamatergic neurons. Bottom: Quantification of center time, total distance and entries to center area (n=6). Student’s t test (two-tailed, unpaired) was applied. ns, not significant; *p < 0.05. Error bars, SEM. (R) Top: Trajectory of mice in open-field chambers with food pellets placed in the center before and after the depletion of LHA glutamatergic neurons. Bottom: Quantification of entries to center area, total distance and number of food pellets consumed by mice in open-field chambers with food pellets placed in the center (n=6). Student’s t test (two-tailed, unpaired) was applied. *p < 0.05. Error bars, SEM.

Further kernel density estimation (KDE) of trans-synaptically retrogradely labeled neurons indicated that the highest density of labeled neurons was located in the caudal region of LHA (cLHA, Figure 1E), which contains both excitatory and inhibitory neurons^32^. Further experiments on optogenetic stimulation of *Vglut2*-ChR2 expressing neurons in 10 hypothalamic nuclei in free-moving mice (Figures S1C-S1E), we found that only the activation of LHA *Vglut2^+^* neurons resulted in robust locomotion of the mice, in a frequency- and light intensity-dependent manner (Figures 1F-H). Thus, LHA *Vglut2^+^* neurons represent the main population of hypothalamic neurons that were involved in regulating mouse locomotion.

### LHA glutamatergic neurons regulate motivation-driven locomotion

To further understand how the firing rate of LHA neurons relates to locomotion speed, we performed single-unit recording at the caudal part of the LHA while mice were walking on a linear track. A total of 156 neurons were recorded from 7 mice. Figure 1I shows an example neuron with a positive correlation with the speed. Analysis revealed a correlation between the firing rate and the speed, using a speed filter previously described ^7^, indicating that neuronal discharge relates to speed changes (Spearman correlation *P*<0.01; *n*=37, median correlation 0.42) (Figure 1J). Subsequently, we applied two widely recognized criteria to categorize these neurons into putative principal cells, inferred to be glutamatergic neurons, and putative interneurons, believed to be GABAergic neurons ^33,34^. Out of these 37 neurons, we found that 24 exhibited characteristics of putative glutamatergic neurons, 3 exhibited characteristics of GABAergic neurons, and the remaining 10 were indistinguishable. (Figure S1F). Moreover, a subpopulation of neurons was found to increase their firing rate prior to the onset of locomotion (Figure 1K). The LHA is known to be critical for regulating various locomotion-dependent physiological and behavioral functions, including predatory attack, evasion, and other motivated behaviors ^9,35,36^. This functional diversity may arise from the heterogeneous cell populations and complex cytoarchitecture within the region. To investigate the association of excitatory versus inhibitory LHA neurons with motivation, we applied fiber photometry to record Ca^2+^ signals of LHA neurons during food seeking. We injected AAV-DIO-GCaMP6s into the LHA of *Vglut2*-Cre or *Vgat-Cre* mice, allowing for the expression of GCaMP6s specifically in either excitatory or inhibitory LHA neurons, thereby facilitating fiber photometry imaging. We observed an increase of Ca^2+^ signals of excitatory neurons that was time locked to the onset of locomotion, with elevated activity during food seeking behavior when approaching the food. However, these signals decreased during the consumption period (Figures 1L and 1M). LHA inhibitory neurons, on the other hand, did not respond to the onset of locomotion and food-seeking behavior; but increased their Ca^2+^ activities during food consumption (Figures S1G and S1H). Meanwhile, in open field test, optical stimulation of LHA *Vgat^+^* cannot induce locomotion in freely-moving mice (Figure S1I).

Next, to further explore the role of LHA *Vglut2^+^* neurons in basic locomotor function and motivated locomotion (e.g., food-seeking), we utilized AAV-DIO-DTR to selectively ablate these neurons. Following diphtheria toxin (DT) administration, no significant changes were observed in basic locomotor functions such as hindlimb protraction, retraction, iliac crest height, or overall activity in an open field setting (Figures 1N-1R), suggesting that the ablation of LHA glutamatergic neurons did not impact basic locomotion. However, in the open field chamber with food pellets located in the center, the locomotor activity was reduced and the mice were less likely to enter the center area, despite the absence of any impact on their appetite measured by food pellet consumption. Together, these results suggest that LHA neurons play a vital role in regulating motivational locomotion.

### LHA neurons regulate locomotion via projections to PnO and MSDB

We next explored the circuit mechanism underlying the function of LHA projection in locomotion control. We searched in a single-neuron projectome dataset for mouse LHA for the major brain regions receiving axon projections from LHA neurons. This dataset consists of single-axon tracing of 916 sparsely labelled neurons expressing various neuropeptides (Figures S2A). To classify the pattern of axon projection patterns of these neurons, we calculated similarity scores based on the shortest distances between neuron pairs. We then performed hierarchical clustering of the similarity matrix of all neuron pairs using Ward’s linkage. Using this approach, we categorized lateral hypothalamus neurons into 4 main clusters and 7 projectome-defined subtypes (Figures 2A-2B, and S2B), with subtypes 1 and 2 in the rostral projecting cluster (Cluster 1, ∼ 16% of the total neurons), subtype 3 in the local projecting cluster (Cluster 2, ∼ 13% of the total neurons), subtypes 4 and 5 in the caudal projecting cluster (Cluster 3, ∼ 46% of the total neurons), and subtypes 6 and 7 in the long-distance projecting cluster (Cluster 4, ∼ 25% of the total neurons).

**Figure 2.**
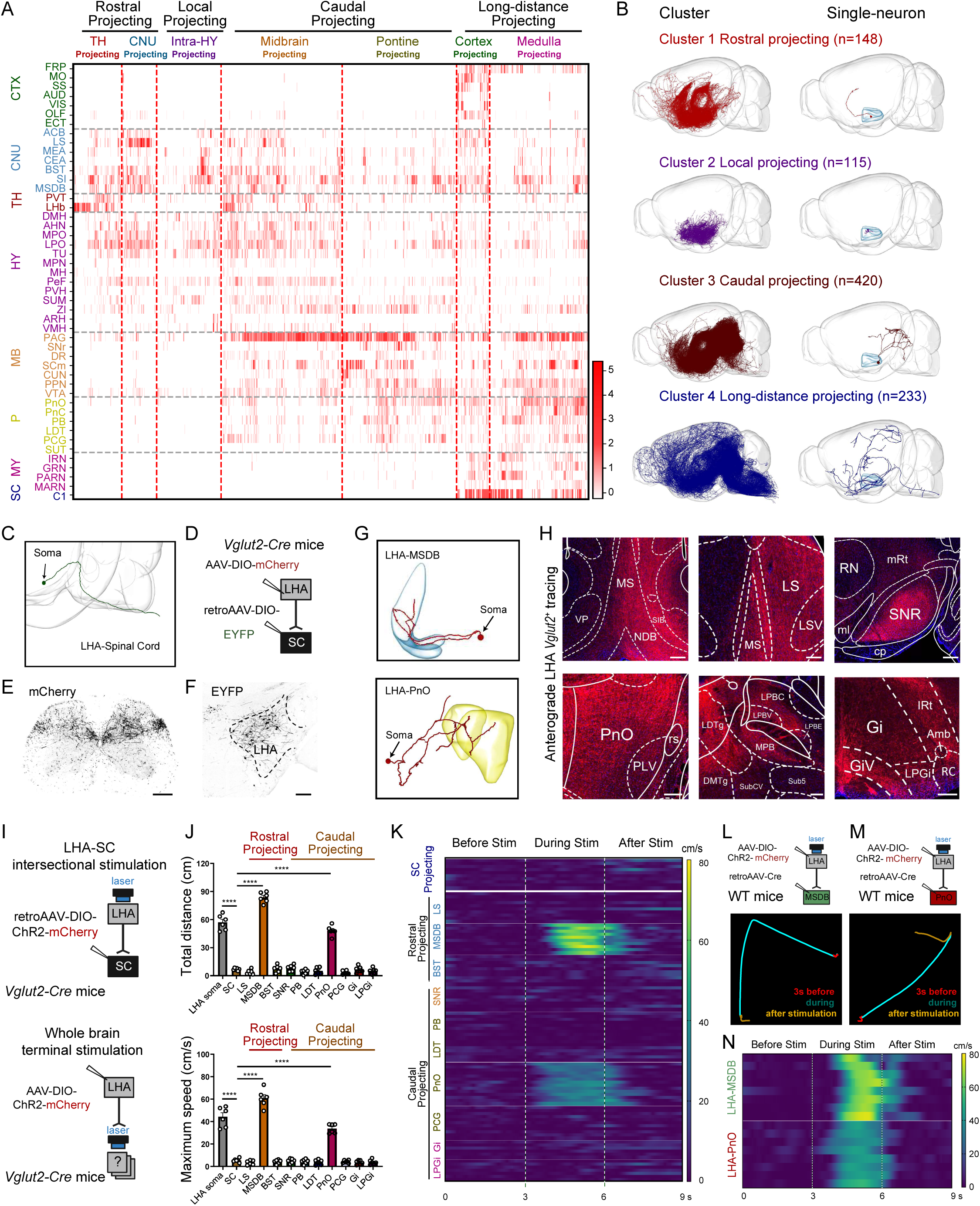
Single-neuron projectome-guided analysis of neural circuits underlying LHA circuit in locomotion control. (A) Projection strength of 916 LHA neurons arranged according to the clustering assignment. Each column represents a neuron. Each row represents a brain region, and the heatmap colors indicate projection strength. (B) Sagittal views of the morphology of all neurons (left) and the example neuron (right) of four classes of LHA neurons. (C) Sagittal views of the morphology of a spinal projecting LHA neuron. (D) Strategy for viral injection to demonstrate the existence of LHA-spinal cord circuit. (E) LHA excitatory projections in lumbar spinal cord. Scale bar, 200 μm. (F) Top: Strategy for retrograde labeling of glutamatergic LHA neurons from lumbar spinal cord. Bottom: Representative image showing spinal cord-projecting LHA glutamatergic neurons. Scale bar, 200 μm. (G) The morphology of representative neurons in the LHA projecting to MSDB (top) and PnO (bottom). (H) Representative images showing different downstream regions of LHA *Vglut2*^+^ neurons from anterograde axonal AAV-DIO-ChR2-mCherry tracing. Scale bar, 200 μm. (I) Strategy for activation of spinal cord-projecting LHA glutamatergic neurons (left) and terminal stimulation of downstream brain regions (right). (J) Quantification of total distance and maximum speed corresponding to optogenetic activation. One-way ANOVA, followed by Bonferroni post hoc test. n = 3 attempts per mouse; n = 6 mice per group. ****p < 0.0001. Error bars, SEM. (K) Heatmap illustrating speed versus time during optical stimulation of LHA neurons projecting to spinal cord and other downstream brain regions. (L) Top: Strategy for activating MSDB projecting LHA neurons. Bottom: Centre of body mass trajectories of single trials in open field arena during 3-s time windows: stationary phase (red), stimulation phase (cyan) and after stimulation offset (orange). (M) Top: Strategy for activating PnO projecting LHA neurons. Bottom: Centre of body mass trajectories of single trials in open field arena during 3-s time windows: stationary phase (Red), stimulation phase (cyan) and after stimulation offset (orange). (N) Heatmap illustrating speed versus time during optical stimulation of MSDB and PnO-projecting LHA neurons.

Among neurons grouped into subtypes 1 and 2 of the rostral projecting cluster (Cluster 1), most displayed targeting preferences for thalamus (TH) and cerebral nuclei (CNU), respectively. Notably, a distinct subgroup within Subtype 1 (TH-projecting) specifically projected to the lateral habenula (LHb). Subtype 2 exhibited strong projections to the cerebral nuclei (CNU), including lateral septal nucleus (LS), medial amygdalar nucleus (MEA), bed nuclei of the stria terminalis (BST), substantia innominate (SI) and medial septal nucleus and MSDB. Cluster 2 consisted of subtype 3, primarily projecting within the hypothalamus with short axons. By comparison, the midbrain-projecting subtypes 4 in cluster 3 projected with varying strengths to the periaqueductal gray (PAG) and superior colliculus (SCm) in the midbrain. Neurons in subtype 5 within cluster 3 primarily terminated their axons within the pontine, including PnO, parabrachial nucleus (PB), laterodorsal tegmental nucleus (LDT), and pontine central gray (PCG). Cluster 4 neurons had broader projections over longer distances, with subtype 6 projecting specifically to the cortex and the medulla, while subtype 7 was more medulla-specific. Among cluster 4 neurons, nearly half (111 out of 233) of the neurons had direct projections to the spinal cord (Figure 2A).

Given that optogenetic activation of LHA glutamatergic neurons elicited hindlimb locomotion (Figure 2C), we examined whether those neurons that projected directly to the lumbar spinal cord circuits, might generate rhythmic motor patterns. Results from anterograde tracing demonstrated that LHA glutamatergic neurons projected robustly to the dorsal part of lumbar spinal cord (Figures 2D and 2E). Retrograde labeling also demonstrated the existence of spinal-projecting LHA glutamatergic neurons (Figure 2F). To explore the function of the LHA-spinal cord pathway, we expressed ChR2 in excitatory spinal cord-projecting LHA neurons by injecting retrogradely transported AAV-expressing Cre-dependent ChR2 into the spinal cord and implanting optical fibers in the LHA of *Vglut2*-Cre mice. We found that optogenetic stimulation of these spinal cord-projecting LHA neurons did not elicit locomotor behaviors (Figures 2I-K). This indicates that an indirect pathway from LHA to spinal cord might be involved.

Based on our hierarchical clustering analysis, anterograde tracing and previous research in motor control ^1,4–6,37,38^, we selected 3 potential downstream brain regions that receive projections from the rostral LHA neurons, including the LS, MSDB, and BST, as well as 7 downstream brain regions that receive projections from the caudal LHA neurons, including the substantia nigra reticular part (SNr), PB, LDT, PnO, PCG, gigantocellular reticular nucleus (Gi), and lateral paragigantocellular nucleus (LPGi), to determine their roles in locomotion (Figures 2G, 2H and S2C). This was achieved by injecting AAVs carrying Cre-dependent ChR2-mCherry into the LHA of *Vglut2*-Cre mice and stimulating the axonal terminals of LHA neurons in these downstream brain regions across the brain (Figures 2I). Optogenetic activation of axon terminals of LHA neurons in these different regions produced varying effects on the mouse locomotion. Notably, activation of LHA neuron projection to the PnO or MSDB could initiate locomotion, whereas MSDB activation led to high-speed locomotion, in contrast to the more moderate responses elicited by PnO activation (Figures 2I-2K).

To further validate the involvement of the LHA-PnO and LHA-MSDB projections in locomotion, we injected retrogradely transported AAVs carrying Cre into the PnO and MSDB, respectively, while expressing Cre-dependent ChR2 in the LHA of wild-type mice. Stimulation of the cell bodies of LHA neurons projecting to PnO or MSDB successfully induced locomotion in freely moving mice (Figures 2L-2N, S2D and S2E). Collectively, these results demonstrate that the projections from the LHA to the PnO or MSDB play a crucial role in regulating locomotion.

### LHA-PnO neurons facilitate motivation-related locomotion without inducing anxiety

Next, we investigated the differential roles of MSDB-projecting and PnO-projecting LHA neurons in regulating mouse locomotion. Our findings demonstrated that optical stimulation of MSDB-projecting LHA neurons elicited high-speed running that resembled escape behavior, a response distinct from that observed following stimulation of LHA-PnO projections. (Figures 2J and 2K). Further open field test and elevated plus-maze test for the stress response of the mice indicated that stimulation of MSDB-projecting LHA neurons resulted in a significant reduction in the time at the center during the open field test and in the time in the open arms during the elevated plus-maze test. In contrast, stimulation of LHA-PnO axon terminals did not elicit any significant stress response (Figures 3A-3C). Through injecting retrogradely transported AAVs carrying Cre into the PnO while expressing Cre-dependent DTR in the LHA of wild-type mice, we found that depletion of PnO-projecting rather than MSDB-projecting LHA neuron terminals impaired motivation-related locomotion, as indicated by the reduced total distance and entries to the center compared to the control group (Figures 3D and 3E). Further examination of the soma distribution of neurons projecting to these two brain areas in single-neuron projectome data showed that somata for these two distinct projections exhibited a spatially distinct anterior-posterior arrangement (Figure 3F and S3A). From connectome data, different projecting clusters exhibited distinct peptidergic signatures (Figures S3B). Interestingly, approximately 48.99% of neurons that project to the spinal cord express Orexin, while subtypes projecting to the PnO and MSDB comprise a heterogeneous population of peptidergic neurons. Axonal collaterals play a crucial role in neuronal communication, allowing for the integration of signals across different regions. Topographic arrangements of individual PnO-projecting neuron revealed the existence of axon collaterals in both PPN and VTA (Ventral tegmental area), which have been reported to play roles in locomotion and motivational circuits, respectively (Figure S3C) ^5,6,9^. Strategy for intersectional viral screening was also performed to identify collateral innervation of PnO-projecting LHA neurons. RetroAAV-Cre was injected in PnO, followed by injection of AAV-DIO-mChery in LHA. Labeling of PnO-projecting LHA neurons revealed collateral axonal processes in the VTA and PPN (Figure S3D). These findings indicate a complex interplay between axonal projection patterns, somatic locations, and neuropeptide expression, which are intrinsic features of specific neuronal populations. Therefore, integrating projectome-defined subtypes with molecular markers and somatic locations is expected to enhance our understanding of the distinct neuronal subtypes involved in diverse hypothalamic functions.

**Figure 3.**
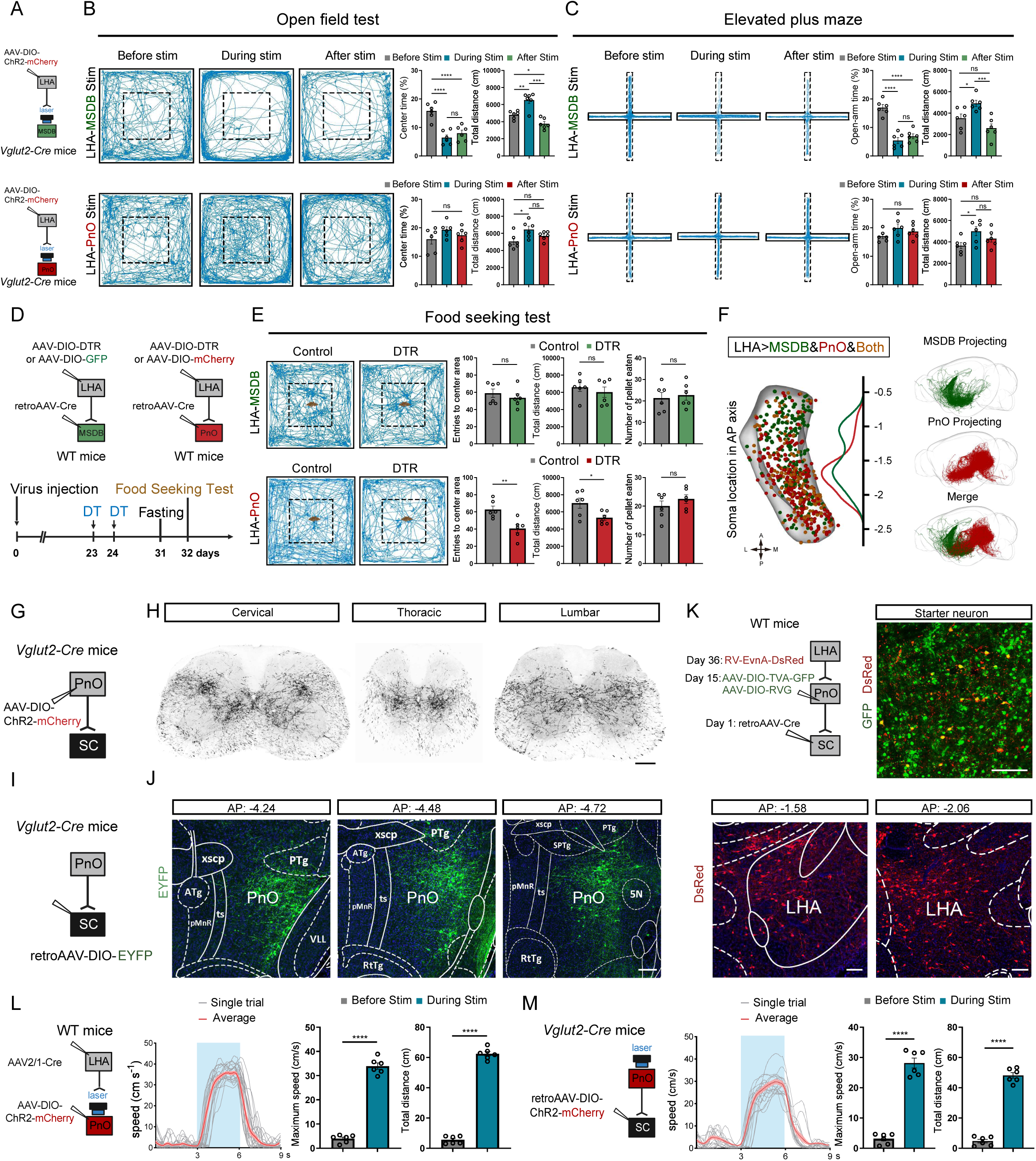
The LHA-PnO-Spinal Cord Pathway is Essential for Relaying Locomotor Signals from the Hypothalamus to the Spinal Cord. (A) Strategy for terminal stimulation of LHA-MSDB and LHA-PnO pathways. (B) Locomotion tracing for an example animal and quantification of center time and total distance of LHA-MSDB (top) or LHA-PnO (bottom) terminal stimulation in the OFT. (C) Locomotion tracing for an example animal and quantification of center time and total distance of LHA-MSDB (top) or LHA-PnO (bottom) terminal stimulation in the EPM. (D) Strategies and timeline summarizing the experiments of food seeking test with or without depletion of MSDB or PnO projecting LHA neurons. (E) Left: Trajectory of mice in open-field chambers with food pellets placed in the center before and after the depletion of MSDB-projecting or PnO-projecting LHA neurons. Right: Quantification of entries to center area, total distance and number of food pellets consumed by mice in open-field chambers with food pellets placed in the center (n=6). Student’s t test (two-tailed, unpaired) was applied. ns, not significant; *p < 0.05; **p < 0.01. Error bars, SEM. (F) Left: Soma location of MSDB (green) and PnO projecting (red) LHA neurons. Right: Arbor reconstructions of MSDB-projecting neurons in subtype 2 and PnO-projecting neurons in subtype 5. (G) Strategy for anterograde labeling of glutamatergic PnO neurons. (H) Representative confocal images showing the expression of mCherry within spinal cord. Scale bar, 200 μm. (I) Strategy for retrograde labeling glutamatergic spinal cord-projecting PnO neurons. (J) Representative confocal images showing the expression of EYFP at indicated positions within PnO. Scale bar, 200 μm. (K) Schematic of three-step monosynaptic rabies virus tracing strategy to demonstrate the circuit of LHA-PnO-spinal cord and trans-synaptically labelled neurons were found in the LHA. Scale bar, 200 μm. (L) Strategy for activation of PnO-projecting LHA neurons. Speed versus time of single trials (grey lines) and the average (red line) of one mouse. Quantification of total distance (left) and maximum speed (right) of mice before and during optical stimulation (n=6). Student’s t test (two-tailed, unpaired) was applied. ****p < 0.0001. Error bars, SEM. (M) Strategy for activation of spinal cord-projecting PnO *Vglut2^+^*neurons. Speed versus time of single trials (grey lines) and the average (red line) of one mouse. Quantification of total distance (left) and maximum speed (right) of mice before and during optical stimulation (n=6). Student’s t test (two-tailed, unpaired) was applied. ****p < 0.0001. Error bars, SEM.

### Activation of the LHA-PnO-Spinal Cord pathway initiates locomotion

Our anterograde tracing experiment showed that PnO comprised a majority of pontine reticulospinal neurons which directly projected to the spinal cord (Figure 3G and 3H). Through retrogradely labeling from the lumbar spinal cord, we found somata of spinal-projecting PnO excitatory neurons were mainly located in the lateral part of PnO (Figure 3I and 3J). To demonstrate the role of PnO as a relay station connecting LHA and lumbar spinal cord, AAVretro-DIO-EYFP was injected into lumbar spinal cord followed by an anterograde tracing virus AAV-DIO-mCherry injected into LHA. Double positive signals were identified in PnO (Figure S3E). Moreover, we performed rabies virus-based three-step monosynaptic retrograde tracing experiments. Specifically, retroAAV-hSyn-Cre was delivered into the lumbar spinal cord, followed by the injection of AAVs expressing Cre-dependent avian-specific retroviral receptor (TVA) and rabies virus glycoprotein (RVG) into the PnO 2 weeks later. RV-EnvA-dsRed was then delivered into PnO and neurons expressing both EGFP and dsRed in PnO represented the “starter cells” in the monosynaptic rabies virus tracing method. Furthermore, trans-synaptically labelled neurons were abundantly found in the LHA (Figures 3K). These results indicate the presence of LHA-PnO-spinal cord pathway. We then expressed anterograde trans-synaptic AAVs (AAV serotype 1) carrying Cre into the LHA ^39–41^, alongside AAV encoding Cre-dependent ChR2 in the PnO to stimulate the LHA-recipient PnO neurons. Optogenetic activation of LHA-recipient PnO neurons produced locomotor effects similar to those observed when stimulating LHA glutamatergic terminals in the PnO, confirming that the LHA-PnO pathway is crucial for locomotion (Figure 3L). To investigate whether PnO-spinal cord pathway directly controls locomotion, we injected retro AAV-DIO-ChR2 into the spinal cord of *Vglut2*-Cre mice. Following ChR2 expression in retrogradely labeled cell bodies in the PnO, optogenetic stimulation reliably induced locomotion in freely-moving mice (Figure 3M). Together, these results indicate that LHA indirectly activate spinal cord neurons through a relay station of PnO to induce locomotion behavior.

### Functional reconstruction of the LHA in a model of spontaneous locomotor recovery following incomplete spinal cord injury

Our results demonstrate that the LHA-PnO-Spinal cord descending circuits play a crucial role in facilitating motivated locomotion in intact mice, leading us to explore their potential involvement in voluntary locomotion following CNS trauma, such as SCI. We first characterized spontaneous locomotor recovery in mice received a lateral hemisection at the thoracic segment 10 (T10) ^21,42^. 3 days post injury, mice displayed a complete loss of stepping ability in the ipsilateral hindlimb, though muscle function in the contralateral hindlimb remained intact. In contrast, in the chronic phase post-SCI, we observed significant locomotor recovery in all subjects, characterized by full-body weight-bearing plantar stepping (Figures S4A and S4B).

Next, we explored whether LHA neurons contribute to the observed recovery. To test this, we used two complementary sets of experiments (Figure 4A). Mice received PRV-labeling from the hindlimb muscle before injury, at the early (1week post-injury) and chronic stage (8 weeks post-injury) after injury. Mice injected with PRV into GS/TA muscle at the early stage exhibited a marked reduction in neuronal labeling across the brain compared to intact mice, indicating disrupted descending pathways to the locomotor circuits below the lesion. In contrast, mice injected with the virus 8 weeks post-injury demonstrated a significant increase of labeled neurons in LHA, suggesting substantial brain reorganization or enhancement of neuronal connections over time.

**Figure 4.**
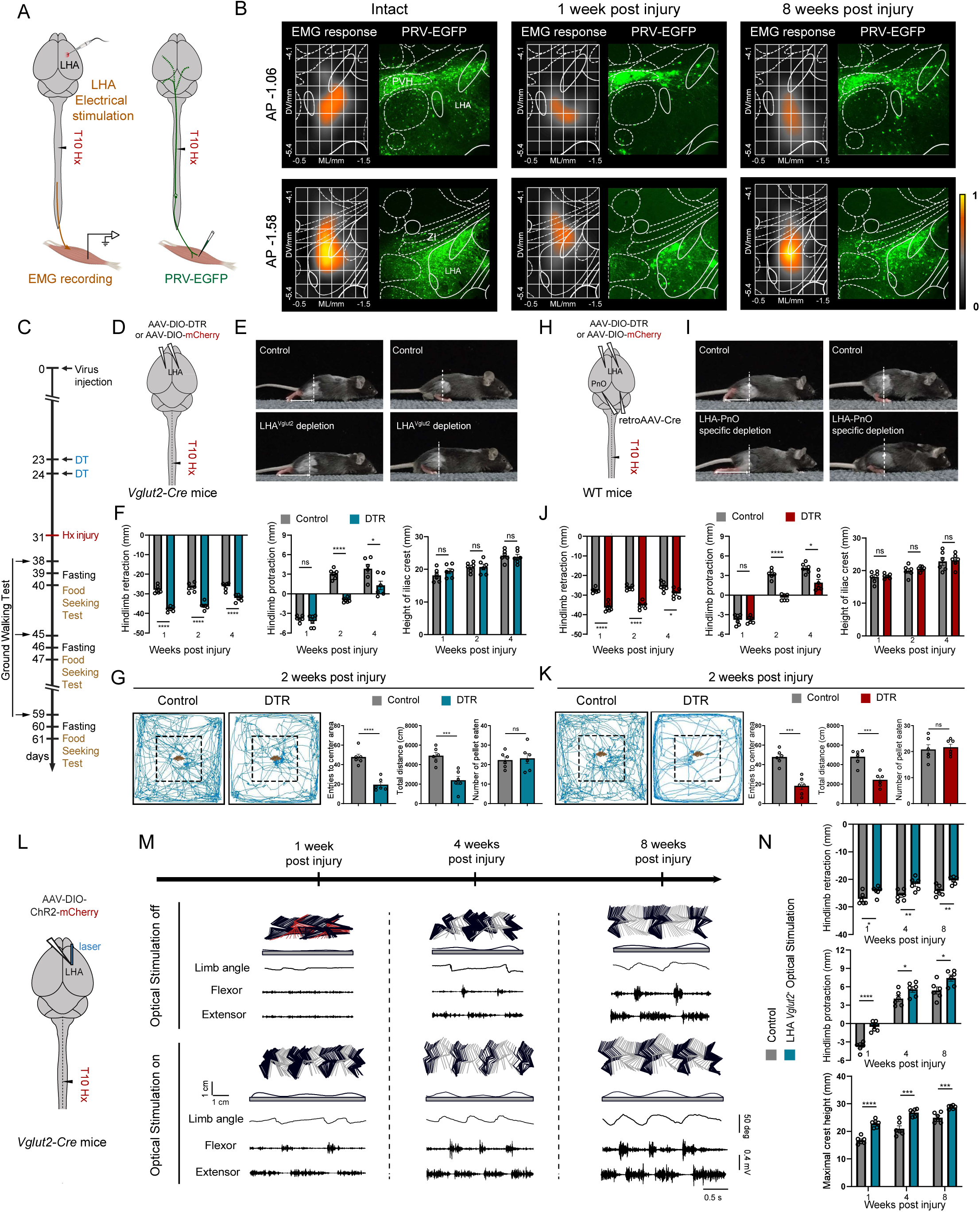
Functional reorganization of LHA-PnO circuits contributes to spontaneous locomotor recovery. (A) Strategy for EMG recording of LHA electrical stimulation and trans-synaptic retrograde labeling of hindlimb muscle related neurons. (B) LHA mapping using electrical stimulation (left) and trans-synaptic PRV tracing (right) before injury, at the acute stage (1w post-injury) or at the chronic stage (8w post-injury). Scale bar, 200 μm. (C) Timeline for locomotor function test and food seeking test after the conditional genetic ablation of LHA *Vglut2^+^* neurons or PnO-projecting LHA neurons in injured mice. (D) Strategy for depletion of LHA *Vglut2^+^* neurons in T10 hemisection injury mouse. (E) Example photographs of mice in ground walking, with or without depletion of LHA glutamatergic neurons after lateral hemisection SCI. (F) Quantification of retraction, protraction and maximum crest height of mice at 1, 2 and 4 wk post T10 lateral hemisection (n=6). Two-way repeated-measure ANOVA, followed by post hoc Bonferroni correction. ns, not significant; *p < 0.05; ****p < 0.0001. Error bars, SEM. (G) Left: Trajectory of mice in open-field chambers with food pellets placed in the center with or without depletion of LHA glutamatergic neurons after hemisection injury mice. Right: Quantification of entries to center area, total distance and number of food pellets consumed by mice in open-field chambers with food pellets placed in the center (n=6). Student’s t test (two-tailed, unpaired) was applied. ns, not significant; ***p < 0.001; ****p < 0.0001. Error bars, SEM. (H) Strategy for depletion of PnO-projecting LHA neurons in T10 hemisection injury mouse. (I) Example photographs of mice in ground walking, with or without depletion of PnO-projecting LHA neurons after lateral hemisection SCI. (J) Quantification of retraction, protraction and maximum crest height of mice at 1, 2 and 4 wk post T10 lateral hemisection (n=6). Two-way repeated-measure ANOVA, followed by post hoc Bonferroni correction. ns, not significant; *p < 0.05; ****p < 0.0001. Error bars, SEM. (K) Left: Trajectory of mice in open-field chambers with food pellets placed in the center with or without depletion of PnO-projecting LHA neurons after hemisection injury mice. Right: Quantification of entries to center area, total distance and number of food pellets consumed by mice in open-field chambers with food pellets placed in the center (n=6). Student’s t test (two-tailed, unpaired) was applied. ns, not significant; ***p < 0.001. Error bars, SEM. (L) Strategy for activating glutamatergic LHA neurons. (M) Representative stick diagram decomposition of leg movements, oscillation of the whole limb (virtual limb linking the hip to the toe) and EMG activity of ankle muscles recorded at 1, 4 and 8 wk post T10 lateral hemisection with and without optical stimulation of LHA *Vglut2^+^*neurons. (N) Quantification of retraction, protraction and maximum crest height of mice at 1, 4 and 8 wk post T10 lateral hemisection (n=6). Two-way repeated-measure ANOVA, followed by post hoc Bonferroni correction. *p < 0.05; **p < 0.01; ***p < 0.001; ****p < 0.0001. Error bars, SEM.

Additionally, we mapped the LHA regions responsible for triggering TA muscle contraction in intact, acute and chronic phase. In intact mice, electrical stimulation of the LHA induced TA muscle contraction, as recorded by electromyography. At the early stage (1week post-injury), stimulation at the same LHA coordinates failed to produce TA muscle contraction. However, stimulation of the same LHA coordinates successfully restored the ability to trigger TA muscle activation 8 weeks post-SCI (Figure 4B). This suggests a re-establishment of functional connections or compensatory neural adaptations within the LHA over time following SCI.

### LHA-PnO circuit facilitates spontaneous locomotor recovery after SCI

Given the reorganization of LHA during spontaneous recovery after incomplete SCI, we evaluated the necessity of LHA glutamatergic neurons for functional restoration. To assess this, we measured basic over-ground locomotion and motivational vigor in fasting-triggered food-seeking behaviors after the conditional genetic ablation of LHA glutamatergic neurons in injured mice (Figure 4C). We found that mice with depletion of LHA *Vglut2^+^*neurons exhibited impaired recovery of locomotor function characterized by pronounced paw dragging and impaired protraction of the denervated hindlimb (Figures 4D-F). The impaired locomotor recovery was associated with decreased motivation for food seeking indicated by decreased total distance and entries to the center area (Figures 4G and S4D).

To specifically deplete PnO-projecting LHA neurons, we injected retrogradely transported AAVs carrying Cre into the PnO while expressing Cre-dependent DTR in the LHA of wild-type mice. Genetic ablation of PnO-projecting LHA neurons significantly impaired functional recovery during over-ground locomotion evidenced by pronounced paw dragging and impaired protraction of the denervated hindlimb (Figures 4H-J). Meanwhile, impaired functional recovery was associated with decreased motivational locomotion after SCI (Figures 4K and S4E). These findings suggest that motivational locomotion encoded by PnO-projecting LHA neurons was essential for natural repair after incomplete SCI.

Subsequently, we asked if activating LHA neurons can promote locomotor recovery at different phases after SCI. To do this, we injected AAVs carrying Cre-dependent ChR2-mCherry into LHA of *Vglut2*-Cre mice. After SCI, mice underwent optogenetic stimulation at 1-, 4- and 8-weeks post injury (Figure 4L). All tested mice demonstrated significantly improved hindlimb locomotion during light stimulation. Detailed kinematic analysis revealed increased weight support and enhanced hindlimb stepping ability (Figures 4M and 4N).

### Acute LHA electrical stimulation enables stepping in paralyzed mice after stagger SCI

We next investigated whether more clinically relevant deep brain stimulation of the LHA could improve locomotion in mice with severe SCI. Our results indicated that stimulation of the LHA-MSDB pathway could induce significant anxiety-like behaviors (Figures 3H and 3I) that were detrimental to recovery. Therefore, we aimed to selectively stimulate the LHA-PnO pathway by adjusting the locus of stimulation within the LHA during DBS.

Based on the somatic distribution of LHA neurons projecting to the PnO and MSDB, we identified that the cell bodies of LHA neurons projecting to the PnO are primarily located in the caudal region of the LHA (Figure 3L), whereas those projecting to the MSDB are predominantly situated in the rostral region. We injected AAVs carrying GCamp6s into both the PnO and MSDB and recorded the activation of these downstream brain regions during stimulation of different locations in the anterior-posterior axis of the LHA. Results showed that stimulating the anterior part of the LHA (−1.0 mm from the bregma) significantly activated the MSDB, while stimulation of the posterior LHA (−1.5 mm from the bregma) resulted in a higher amplitude activation of the PnO (Figures 5A-F). Moreover, electrical stimulation in caudal part within LHA induced locomotion immediately in intact mice (Figures 5G-K). These results guided us to apply LHA DBS in a specific location to facilitate functional recovery following severe SCI.

**Figure 5.**
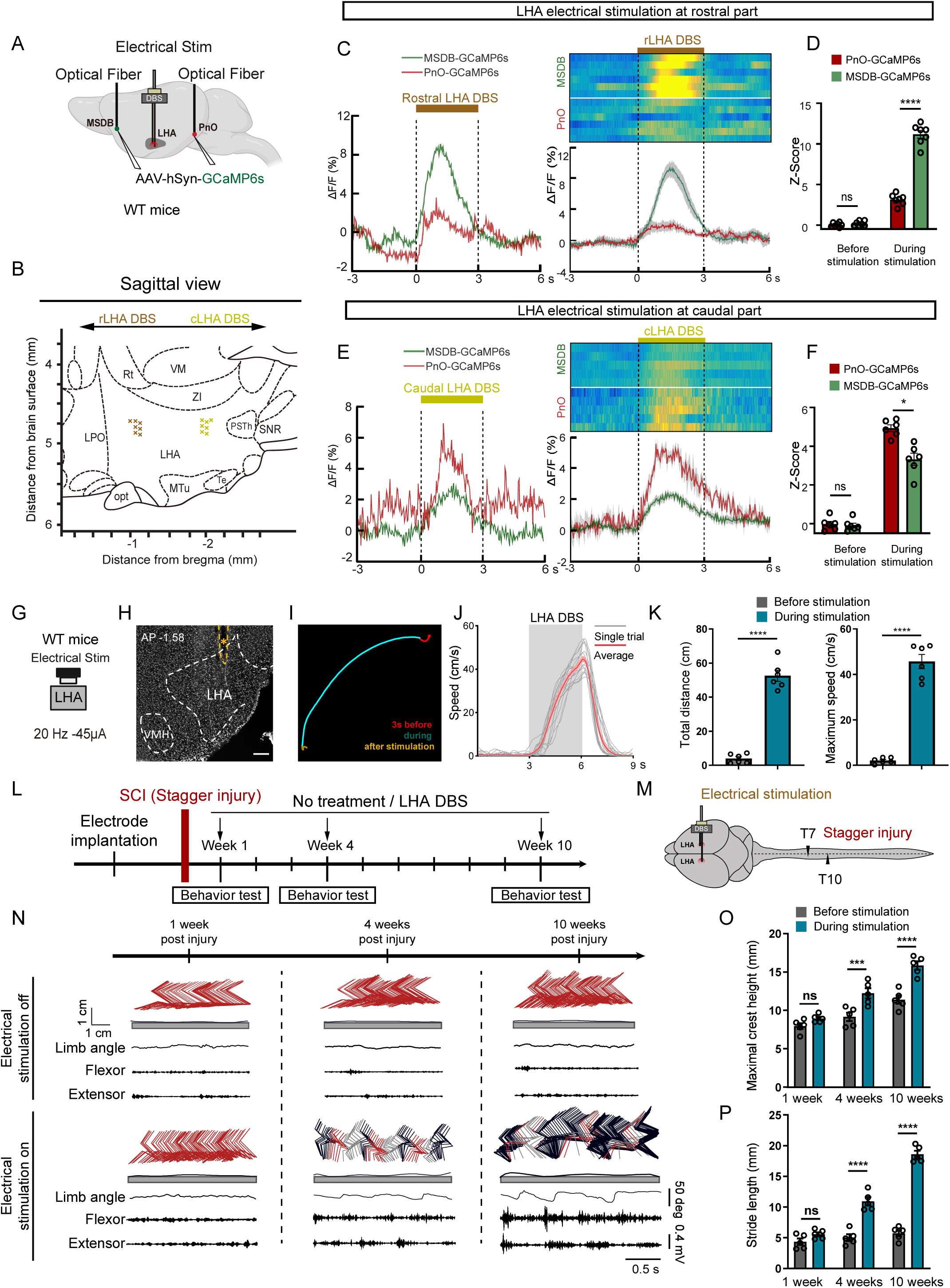
Optimized LHA DBS acutely enables stepping in paralyzed mice after chronic stagger SCI. (A) Strategy for the simultaneous recording of responses from PnO and MSDB to stimulation applied on the LHA. (B) Sagittal view of the locus of electrical stimulation on the rostral and caudal part of LHA. (C-D) The results of electrical stimulation at rostral part of LHA. The calcium signals of an example trial (left) and all trials (right) are shown (C). MSDB was predominantly activated when the rostral part of LHA was stimulated (D, p<0.001). (E-F) The results of electrical stimulation at the caudal part of LHA. The calcium signals of an example trial (left) and all trials (right) are shown (E). PnO was activated to a greater extent than PnO when the caudal part of LHA was stimulated (F, p<0.05). (G) Strategy for electrical stimulation of LHA in intact mice. (H) Coronal view of post-hoc anatomical evaluation of the electrode placement in LHA, in the vicinity of the DBS electrode implantation site. Scale bar, 200 μm. (I) Centre of body mass trajectories of a single trial in open field arena during 3-s time windows: stationary phase (red), electrical stimulation phase (cyan) and after stimulation offset (orange). (J) Speed versus time of single trials (grey lines) and the average (red line) of one mouse. (K) Quantification of total distance (left) and maximum speed (right) of mice before and during electrical stimulation of LHA (n=6). Student’s t test (two-tailed, unpaired) was applied. ****p < 0.0001. Error bars, SEM. (L-M) Timeline and scheme summarizing the experiments. A severe SCI model of staggered lateral hemisections at T7 and T10. (N) Representative leg kinematics and hindlimb EMG data from SCI mice in the acute injury (7 DPI), and sub-chronic (28DPI) and chronic injury stages (70DPI). Scale bar, 0.4mv, 0.5s. Quantification of maximal crest height (O) and stride length (P) of mice at 1w, 4w and 8w post-hemisection injury (n = 6 and n = 5 respectively). Student’s t test (two-tailed, unpaired). *p < 0.05; **p < 0.01. Error bars, SEM.

Given the enhancement of locomotor recovery via activating the LHA-PnO descending pathway in lateral hemisection SCI mice, we questioned if similar manipulation could restore function in severe SCI models. Therefore, we employed a double lateral hemisection SCI model at T7 and T10 levels, creating a more challenging scenario by eliminating brain-derived innervation below the second injury site while maintaining a spared tissue bridge between lesions to relay descending signals ^17,43^. We conducted a timeline experiment to assess the effects of LHA-DBS on locomotor function, evaluating animals with and without LHA-DBS at post-injury weeks 1, 4, and 10, and analyzing immediate effects on hindlimb stepping (Figures 5L and 5M). Initially, no hindlimb responses were observed with LHA-DBS at 1week post-injury. However, by 4 weeks after injury, animals exhibited significantly improved walking ability during LHA-DBS. Continuous DBS promoted pronounced locomotor improvements, as indicated by significant increases in maximum iliac crest height and stride length. 10 weeks post-injury, mice exhibited better locomotor functional recovery with LHA-DBS (Figures 5N-P).

Subsequently, we aim to elucidate the mechanism by which LHA-DBS facilitates functional improvement in descending input-dependent movement. The activation of thoracic long-distance projecting neurons ^44^ through spared relay pathways, such as axonal sprouting of the reticulospinal tract ^21,22^, plays a pivotal role in the restoration of locomotor function following SCI. Consequently, the mechanisms underlying LHA-DBS are likely dependent on the activation of Zfhx3 neurons, which are predominantly found among long-distance projecting neurons ^45^. Target DBS at cLHA led to a significant increase in cFos-positive neurons, as well as c-Fos and Zfhx3 double-positive neurons, in the inter-lesion spinal cord at 4- and 10-weeks post-injury, compared to the early stage following SCI (Figures S5A-S5C).

However, behavioral improvement was not maintained after the cessation of stimulation, necessitating further investigation into the potential of LHA-DBS for facilitating long-term locomotor functional recovery after SCI.

### Brain-controlled LHA-DBS enhances locomotor recovery after sever SCI

Previous findings have demonstrated that the delivery of DBS triggers stress responses in animals ^27^ and elevated stress levels were also observed under the condition of stochastic LHA-DBS despite stimulating specific locus at cLHA. To alleviate stress levels, we applied unsupervised learning algorithms to trigger LHA-DBS.

The intention of locomotion can be decoded from the multi-unit activity (MUA) in the motor cortex’s hindlimb region (M1HL) ^27^. Therefore, decoding this locomotor initiation information from the motor cortex could enable transient stimulation of the LHA via a brain-computer interface (BCI)-DBS system. To explore this, mice were outfitted with a 32-channel micro-wire array electrode implanted in M1 to record MUA and decode locomotor intentions. MUA firing rate increased prior to the initiation of hindlimb movements, observable in both intact and staggered SCI mice (Figures 6A and 6B). Under conditions of brain-controlled electrical stimulation, LHA-DBS did not provoke stress-like responses in either healthy or injured mice (Figures 6C-6F).

**Figure 6.**
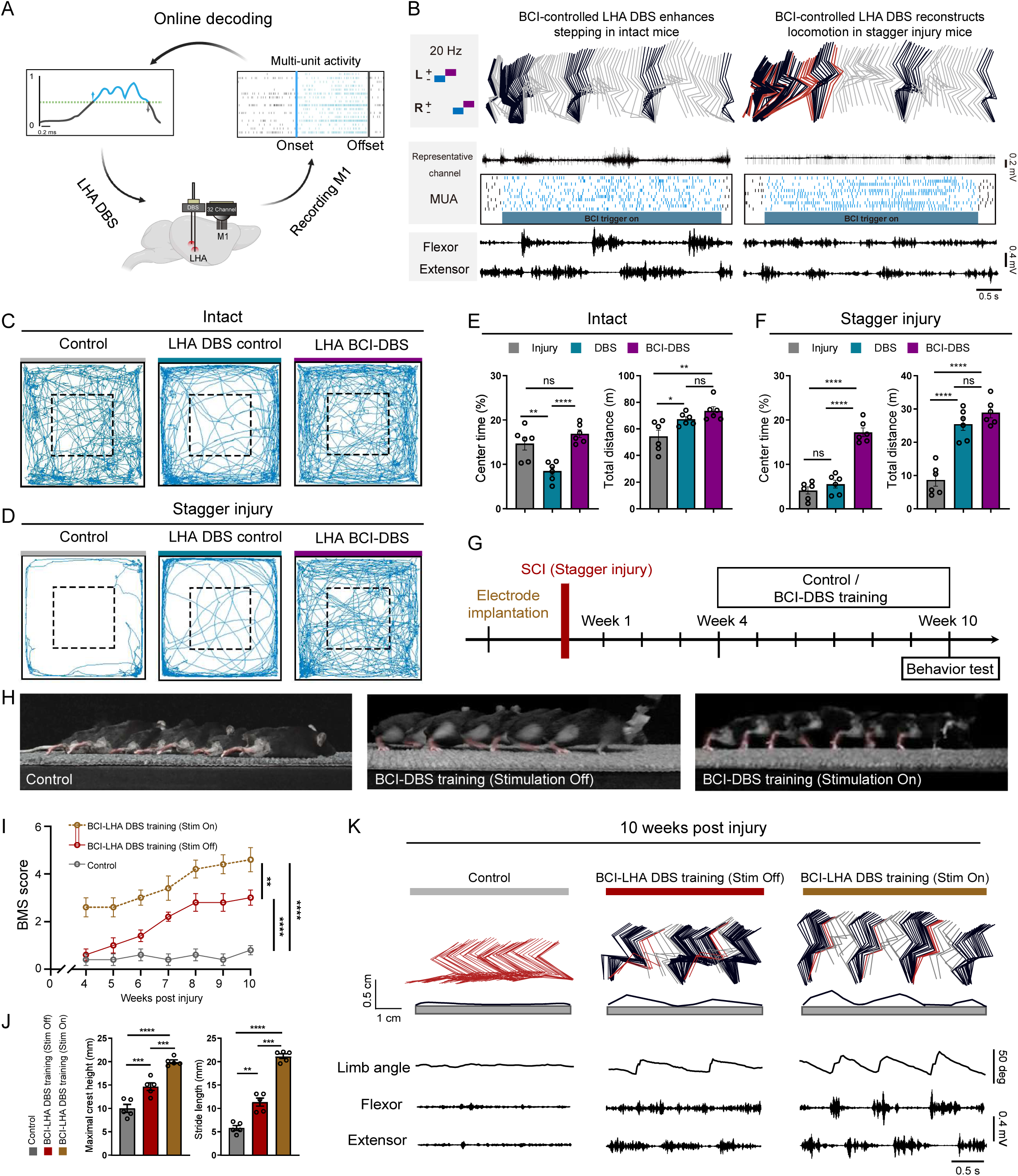
Brain-Controlled LHA-DBS Enhances Locomotor Recovery after sever SCI. (A) The schematic diagram of LHA DBS triggered by BCI. A 4*8 array was used to record neural signals from M1. The multi-unit activity (MUA) was processed online by an unsupervised algorithm. DBS application and intensity depended on the neural activities. (B) Representative leg kinematics, neural activities, and hindlimb EMG data of healthy mice (left) and staggered mice (right). Scale bar of representative neuronal active channel, 0.2 mV. Scale bar of EMG data, 0.4mV, 0.5s. (C) Trajectory of intact mice in open-field chambers without DBS (left), with irregular DBS (middle), and with brain-controlled DBS (right). (D) Trajectory of mice after SCI (stagger injury) in open-field chambers without DBS (left), with irregular DBS (middle), and with brain-controlled DBS (right). (E) Quantification of the proportion of center time (left) and total distance (right) in intact mice, comparing no DBS to irregular or brain-controlled DBS. Student’s t test (two-tailed, unpaired). *p < 0.05; **p < 0.01. Error bars, SEM. (F) Quantification of the proportion of center time (left) and total distance (right) in post-SCI (stagger injury) mice, comparing no DBS to irregular or brain-controlled DBS. Student’s t test (two-tailed, unpaired). *p < 0.05; **p < 0.01. Error bars, SEM. (G) Timeline summarizing the experiments about the effect of LHA brain-controlled DBS in the recovery process of mice following stagger SCI. (H) Chronophotography of mice illustrates the recovery process following a stagger injury. (I) Representative leg kinematics and hindlimb EMG data from mice without training (left), trained without stimulation (middle) and trained with stimulation (right) after stagger injury. (J) Quantification of maximal crest height (left) and stride length (right) of mice at 70 DPI without training, trained without DBS or trained with DBS. (K) BMS test performance in control, BCI-DBS training without stimulation and BCI-DBS training with stimulation groups. n=LJ5 per group, two-way ANOVA-RM with Bonferroni post hoc correction. **p < 0.01, ****p < 0.0001. Error bars, SEM.

These findings suggest that a BCI-DBS system could serve as a viable long-term therapeutic approach. To test this, staggered SCI mice underwent daily training sessions for 6 weeks, with brain-controlled DBS starting 4 weeks post-injury (Figure 6G). In contrast to the paralyzed control group, long-term BCI-DBS training resulted in significant functional recovery, including restored hindlimb stepping ability and increased BMS scores. Notably, after prolonged training, mice displayed improved locomotor performance during ground walking with stimulation (Figures 6H-I). Kinematic analyses revealed substantial improvements in locomotion compared to control groups. During ground walking, mice subjected to long-term BCI-DBS training were able to partially support their body weight, showed significantly increased hindlimb oscillation, and exhibited muscle activity as confirmed by electromyogram (EMG) recordings. Additionally, those BCI-DBS training mice demonstrated enhanced locomotor performance when stimulation was applied (Figures 6J-K). Furthermore, long-term BCI-DBS training led to a significant increase in cfos-positive and cfos/Zfhx3 double-positive neurons in the inter-lesion spinal cord (Figures S6A-S6C). Collectively, these results indicate that long-term brain-controlled LHA-DBS training effectively promotes locomotor recovery.

## DISCUSSION

The lateral hypothalamus, an evolutionarily conserved brain region, receives significant attention in the contexts of emotion, energy metabolism, and innate behaviors. Nonetheless, the elucidation of its function in the transmission of initiation commands to the spinal cord, as well as its prospective role in therapeutic interventions for disease, remains an area of ongoing investigation ^46^. In this study, we identified a population of glutamatergic neurons in the caudal LHA (cLHA) that transmit excitatory signals to pontine reticulospinal neurons in the PnO, which in turn project to the spinal cord to initiate locomotion. We demonstrated that those PnO-projecting LHA *Vglut2^+^* neurons play a crucial role in motivated locomotion during food seeking behavior. Additionally, these neurons contribute to various aspects of motor recovery following incomplete spinal cord injury (SCI). Our findings underscore the importance of LHA circuits in transmitting locomotor commands to the spinal cord, advance our understanding of how supraspinal circuits facilitate functional recovery after SCI, and highlight a promising target for therapeutic interventions.

### Glutamatergic neurons in the cLHA initiate locomotion via pontine reticulospinal projections

Despite the lack of a fully defined anatomical boundary for the SLR, previous studies have identified regions within the LHA or its surrounding areas ^47,48^ can induce locomotion independent of the MLR ^2,49^. This independence has led to speculation about the SLR’s specific role in locomotor control. By delivering the PRV virus to the hindlimb muscles, our results revealed an extremely densely labeled region in the caudal part of the LHA. Further findings indicate that activating *Vglut2^+^* neurons, rather than *Vgat^+^* neurons, in this area can induce robust locomotion. LHA is a particularly complex region, containing heterogeneous populations of peptidergic neurons with widespread projections. This complexity has made it difficult to fully decipher the LHA’s role in locomotor control ^32,50^. Recent studies have provided insights into specific neuronal subtypes within the LHA that are involved in regulating movement. For example, orexin neurons respond to movement on a millisecond timescale ^51^, and LHA neurons projecting to the PAG ^35^ or VTA ^36^ mediate evasion or defensive behavior, respectively. However, the exact pathways by which the LHA regulates locomotion remain incompletely understood. We found that the PnO-projecting glutamatergic LHA neurons can reliably induce forward locomotion, and the projections from PnO to SC serve as a relay downstream pathway for this behavior. In conclusion, it’s noteworthy that we discovered an LH-PnO-SC excitatory circuit that can control locomotion without inducing negative emotional effects.

LHA has extensive projections, the organization of which remains challenging to fully elucidate. In this study, high-resolution fMOST analysis classified its single-neuron projections into four major categories. Notably, a group of long-projecting neurons to the spinal cord, with approximately 48.99% expressing orexin (FigureS3), was identified. However, activation of these direct spinal cord-projecting neurons in the LHA did not induce locomotion. In contrast, activation of neurons projecting to the PnO in the cLHA, including about 21.58% orexin-positive neurons (Figure S5C), robustly initiated locomotion. The fMOST analysis also revealed intriguing projection patterns, including collateral projections to both the PnO and the PPN (Figure S3D), which play important roles in exploration. However, fewer neurons projected to the CnF, which is involved in escape behaviors. Taken together, these findings suggest that the LHA controls locomotion via pontine reticulospinal projections, functioning through an MLR-independent circuit, but potentially also through a PPN-dependent circuit. Other studies have also suggested that the PnO can transmit a variety of action commands, such as controlling locomotor gait asymmetries or inducing behavioral arrest ^39,52^. These findings highlight the functional diversity of PnO neurons, and future investigation is warranted to clarify the precise relationships between these circuits.

### The LHA circuit in emotional valence processing

The LHA’s ability to integrate arousal, motivational and evasive underscores its role as a central hub for processing of emotional valence ^10,53,54^. Rather than functioning in isolation, the LHA is part of a broader network of brain regions that collectively shapes behavior in response to internal drives and external challenges ^36,55^ or primarily mediates appetitive locomotion ^10,56–58^. Our study demonstrated that LHA *Vglut2^+^* neurons projecting to the PnO play a crucial role in facilitating motivated locomotion, particularly during goal-directed behaviors like food-seeking. Importantly, locomotion was observed without the induction of anxiety or stress-like behaviors, indicating that the LHA-PnO circuit can selectively enhance motor functions associated with motivation without triggering negative emotional states. This finding is particularly interesting because it dissociates the locomotor and emotional functions of the LHA, which has traditionally been closely associated with both motor control and emotional regulation ^36^.

While the LHA-PnO circuit facilitates motivated locomotion, a distinct projection pathway from the LHA to the MSDB appears to be involved in evasion and anxiety-like behaviors (Figures 3H and 3I). Neurons in the rostral LHA primarily project to subcortical (subtype 1) and thalamic areas, such as the lateral septum (LS) and lateral habenula (LHb), suggesting a key role for the LHA in emotional regulation, consistent with past research ^59,60^. Our findings revealed that activation of LHA *Vglut2^+^* neurons projecting to the MSDB (subtype 2) induced anxiety-like behaviors, suggesting that this circuit plays a role in emotional responses to potential threats. Furthermore, intersectional stimulation of the MSDB and LHA resulted in escape-like high-speed movements, reinforcing the idea that this pathway is involved in defensive or evasive behaviors. The LHA-MSDB circuit’s role in anxiety and evasion aligns with previous research showing the MSDB’s involvement in modulating emotional states, particularly those related to fear and anxiety ^61,62^. Our findings extend this knowledge by demonstrating that LHA *Vglut2^+^* neurons projecting to the MSDB can directly influence these emotional responses, likely through their extensive projections to subcortical and thalamic areas (Figure 2A). This circuit’s involvement in both anxiety and escape-like behaviors suggests that it plays a critical role in coordinating negative emotion and motor responses ^37,61,63^.

The dissociation between the LHA-PnO and LHA-MSDB circuits in terms of locomotion and anxiety-like behaviors suggests that different subsets of LHA *Vglut2^+^* neurons are functionally segregated, with some circuits promoting motivated actions and others modulating emotional states. This circuit-leveled dissociation is intriguing, as it suggests that activation of select LHA neurons involved in motor control can promote locomotion in intact and SCI mice without exacerbating negative emotional states, such as anxiety or depression ^60^. This is particularly important in the context of SCI, where emotional disturbances often can hinder the rehabilitation efforts to recover locomotion ^64^.

### Therapeutic potential of DBS and its implications for clinical translation in SCI

By elucidating the circuit mechanisms by which the LHA controls locomotion, we have expanded our understanding of using the DBS application to promote functional recovery after SCI. Our findings show that DBS stimulation of LHA, particularly targeting its glutamatergic neurons, significantly improved locomotor function in paralyzed mice. Guided by viral tracing and high-resolution fMOST analysis, we optimized the stimulation locus, primarily activating the LHA-PnO pathway rather than the LHA-MSDB pathway. This approach facilitated the restoration of locomotor function in SCI mice while minimizing the induction of stress. To maximize the efficacy of DBS, we further developed a brain-controlled DBS system that leverages real-time decoding of motor cortex activity to synchronize LHA stimulation with the intention to walk. This closed-loop system not only facilitated immediate improvements in motor function but also promoted long-term restoration of hindlimb motor function. By aligning LHA stimulation with the animal’s intention to walk, this approach effectively harnesses the brain’s natural drive to move, amplifying the effects of DBS and promoting more targeted rehabilitation. Our study thus represents a significant advancement in the field of neuromodulation for functional SCI recovery, as it allows for more precise and personalized interventions.

### Limitations and future directions

Despite the promising results of our study, several limitations must be acknowledged. First, although we demonstrated that LHA stimulation enhances locomotor recovery in mice, future studies should explore the efficacy of LHA-DBS in more clinically relevant models, such as non-human primates, which may more closely resemble the conditions of SCI patients. Second, the degree to which these findings can be translated to humans remains uncertain. Human spinal cord injuries are often more complex than the experimental models used in this study, involving a wider range of injury patterns, comorbidities, and longer recovery timelines. Further research is needed to determine whether similar mechanisms of motor recovery are applicable to human patients. This will require sophisticated clinical trials and the development of more refined models that better replicate the complexity of human SCI.

Another important area for future investigation involves the potential effects of LHA-DBS on eating behaviors, given the well-established role of the lateral hypothalamus in regulating hunger and satiety. The LHA, particularly its inhibitory neurons, is known to be critical for controlling feeding behavior, with dysregulation in this region linked to conditions such as anorexia or hyperphagia ^65^. The LHA activity indicates hunger and satiety states in humans ^66^. Patients subjected to LHA-DBS significantly altered food motivation, although the specific mechanisms of motor function modulation by the LHA require further investigation. Given this, it is essential to consider the possible unintended consequences of LHA-DBS on eating behaviors, especially in patients who may already be vulnerable to metabolic or psychological disturbances.

In conclusion, our study identifies the lateral hypothalamus (LHA) as a critical driver of locomotor initiation and recovery following spinal cord injury (SCI). By modulating the activity of the pontine reticulospinal tract, the LHA enhances motivated locomotion without triggering stress responses, presenting a novel target for therapeutic intervention. Additionally, our development of a brain-controlled deep brain stimulation (DBS) system underscores the potential of neuromodulation to promote functional recovery after SCI. These findings mark a significant advance in understanding the brain’s role in locomotor recovery and offer a promising strategy for restoring motor function in individuals with severe SCI. Future research should focus on translating these insights into clinical applications, with an ultimate goal of improving outcomes for patients suffering from spinal cord injuries.

## STAR★METHODS

Detailed methods are provided in the online version of this paper and include the following:

- KEY RESOURCES TABLEONTACT FOR REAGENT AND RESOURCE SHARING
- EXPERIMENTA MODEL AND SUBJECT DETAILS

- Mouse Strains
- METHOD DETAILS

- Retrograde trans-synaptic PRV tracing
- Transparency and 3D image acquisition
- Stereotactic surgery
- Single-neuron projectome: data acquisition and analysis
- DTR-mediated cell ablation
- Optical-fiber-based Ca2+ recording in freely behaving mice
- Single unit recording and analysis
- EMG Recording
- Behavioral experiments and kinematic analysis
- Open-field (OF) test Food seeking test and elevated plus maze (EPM)
- SCI Surgical procedures
- Post-surgical treatments and care of the animals
- Close-looped Deep brain stimulation
- Immunohistochemistry and Imaging
- QUANTIFICATION AND STATISTICAL ANALYSIS

## ACKNOWLEDGMENTS

We thank Drs Zhigang He, Philip Williams and Yang Xiang for critically reading the manuscript. We thank the CEBSIT Mouse Brain Mesoscopic Connectome Core Facility for their assistance in data analysis and the Gene Editing Core Facility for their assistance in virus technology services. This work was supported by the National Natural Science Foundation of China (92168105 to Y. Li, 82301563 to Z. Li, 82401616 to S. Yu, 82272715 to J. Chen, 81974335 and 82172426 to W. Cai.) and Biosecurity Research Project (23SWAQ24 to Y. Li and J. Chen). Y. Li acknowledges the support from Shanghai Municipal Science and Technology Major Project (2018SHZDZX05). W. Cai acknowledges the support from Major Basic Research Fund of Jiangsu Province Hospital (TS202402).

## AUTHOR CONTRIBUTIONS

Y.L. of the project. C.J., Z. Lin., Y.Z., Z. Zhao., Z.J., Z. Zheng., Z. Li., S.Y., Y.Q., Y.W., A.S., H.S., and Q.W. performed the experiments and discussed the results. X.X., J.C., B.C., W.C. X.W. and X.S. participated in data analysis, Y.L., C.J. and Z. Lin. prepared the manuscript with input from all authors and all authors were involved in interpretation of experiments and contributed to writing the paper.

