## Supplementary material for "Activation of hypothalamic-pontine-spinal pathway promotes locomotor initiation and functional recovery after spinal cord injury in mice": Methods

**STAR★METHODS**

KEY RESOURCES TABLE

| REAGENT or RESOURCE | SOURCE | IDENTIFIER |
| --- | --- | --- |
| Antibodies | | |
| Chicken anti-GFP | Aves Labs | GFP-1020 |
| Goat anti-cholinergic acetyltransferase (ChAT) | Merck | AB144P |
| Rabbit anti-RFP | Abcam | ab34771 |
| Rabbit anti-cfos | CST | 2250s |
| Donkey anti-chicken 488 | Jackson ImmunoResearch | 703-545-155 |
| Donkey anti-goat 647 | Thermo Fisher | A21447 |
| Donkey anti-rabbit cy3 | Jackson ImmunoResearch | 711-165-152 |
| Virus Strains |  |  |
| PRV-CAG-EGFP | Braincase | BC-PRV-531-Plus |
| AAV2/9-EF1α-DIO-DTR-tdTomato | BrainVTA(Wuhan) Corp., Ltd | PT-8328 |
| AAV2/9-hSyn-DIO-GCaMP6s | BrainVTA(Wuhan) Corp., Ltd | PT-0091 |
| AAV2/9-EF1α-DIO-ChR2-mCherry | BrainVTA(Wuhan) Corp., Ltd | PT-0002 |
| AAV2/2Retro-hSyn-Cre | BrainVTA(Wuhan) Corp., Ltd | PT-0136 |
| AAV2/8-EF1α-DIO-H2B-EGFP-T2A-TVA | BrainVTA(Wuhan) Corp., Ltd | PT-0021 |
| AAV2/8-EF1α-DIO-RVG | BrainVTA(Wuhan) Corp., Ltd | PT-0023 |
| RV-EvnA-DsRed | BrainVTA(Wuhan) Corp., Ltd | R0302 |
| AAV2/1-hSyn-Cre | BrainVTA(Wuhan) Corp., Ltd | PT-0136 |
| AAV2/2Retro-EF1α-DIO- ChR2-EYFP | BrainVTA(Wuhan) Corp., Ltd | PT-0001 |
| Chemicals, peptides, and recombinant proteins | | |
| Diphtheria toxin | Sigma | D0564-1MG |
| Clozapin N-oxide | BrainVTA | CNO-01-1103 |
| Experimental models: organisms/strains | | |
| Mouse/ C57BL/6 | SLAC laboratory Animal, Shanghai |  |
| Mouse/ *Vglut2*-ires-Cre | The Jackson Laboratory | Stock 016963 |
| Mouse/*Vgat*-ires-Cre | The Jackson Laboratory | Stock 016962 |
| Software and algorithms | | |
| MATLAB (v2018a) | Mathworks | <https://www.mathworks.com/> |
| Python (v3.11) | Python | <https://www.python.org/> |
| ImageJ2/Fiji | NIH | <https://imagej.nih.gov/ij/> |
| GraphPad PRISM (v7.0) | GraphPad PRISM | <https://www.graphpad.com:443/> |
| DeepLabCut | Mathis Lab (Mathis et al., 2018) | <http://mackenziemathislab.org/deeplabcut> |

**CONTACT FOR REAGENT AND RESOURCE SHARING**

**EXPERIMENTAL MODEL AND SUBJECT DETAILS**

**Mouse Strains**

The following mouse lines were used in this study: wild-type C57BL/6J (SLAC laboratory Animal, Shanghai), *Vglut2*-ires-Cre mice (The Jackson Laboratory, Stock 016963), *Vgat*-ires-Cre mice (The Jackson Laboratory, Stock 016962). Eight-to twelve-week-old male and female mice were randomly assigned to experimental groups. All mice were housed under consistent conditions with unrestricted access to food and water and maintained on a 12-h light/dark cycle (lights on from 7:00 a.m. to 7:00 p.m.). Before food seeking test, mice were fasted for 24 hours. All animal studies and experimental procedures received approval from the Animal Care and Use Committee of the Institute of Neuroscience, Chinese Academy of Sciences.

**METHOD DETAILS**

**Retrograde trans-synaptic PRV tracing**

In the retrograde trans-synaptic tracing experiments, mice were anesthetized with intraperitoneal injection of Zoletil 50 (30mg/kg) in combination with Xylazine Hydrochloride (40mg/kg) and placed on the surgical table. An incision was made on the left to expose the tibialis anterior muscle. Three injection sites were made at a depth of 0.5-1.0 mm into the muscle through the microliter syringes (Hamilton). At each site, 1 μL PRG-EGFP was injected. Following the PRV injection, the incision was sutured. For histological analysis, mice were perfused at 5.5 days after the PRV injection.

**Transparency and 3D image acquisition**

All harvested samples were post-fixed overnight at 4°C in 4% PFA in PBS. The solution of Benzyl Alcohol and Benzyl Benzoate (BABB) was then used to match the refractive index of the tissue. Cleared samples were imaged in a transverse orientation on a light-sheet microscope. The samples were scanned using the continuous light- sheet scanning method with the included contrast blending algorithm for the 564 nm channels. Cell detection was performed with the newly developed open-source ClearMap software (Renier et al., 2014), tailored for cell nuclei, which used a background subtraction via morphological opening, followed by a sequence of filters, morphological operations, and a 3D peak detection. The color of the bubble corresponds to n^2^ fold change in cell density in HY.

**Stereotactic surgery**

Stereotactic injections into both the brain and spinal cord were performed using a microinjection syringe pump (World Precision Instruments, WPI) and a stereotactic apparatus (RWD Life Science). Prior to surgery, mice were anesthetized by intraperitoneal injection of Zoletil 50 (30mg/kg) in combination with Xylazine Hydrochloride (40mg/kg).

For the stereotactic injections into the brain, the skull of the head-immobilized mouse was exposed and treated with 3% hydrogen peroxide to remove connective tissue from the surface. The skull was then leveled anteroposteriorly between the bregma and lambda, and mediolaterally between the left and right hemispheres. A small hole was drilled in the skull at the precise coordinates corresponding to the target brain region. Virus was delivered to the target brain region at a rate of 50 nL/min using the microinjection syringe pump (WPI). When the injection was completed, the micropipette was held in place for 10 min, then raised up by 0.1 mm following an additional 3 min, and finally withdrawn at a low speed.

The coordinates for different brainstem targets are as follows (zero from Bregma; AP, anterior-posterior; ML, medial-lateral; DV, dorsal-ventral from dura; in mm): LHA, AP -1.58, ML ± 1.0, DV 4.95; PnO, AP -4.3, ML ± 0.85, DV 3.95; MSDB, AP +1.00, ML ± 0.10, DV 3.60; M1, AP -0.7 to -1.2, ML ± 1.0, DV 0.55; Gi, AP -6.0, ML ± 0.35, DV 5.10; SNr, AP -3.2, ML ± 1.15, DV 4.30; LPGi, AP -6.90, ML ± 1.10, DV 5.00; MLR, AP -4.80, ML ± 1.19, DV 2.70; LDT, AP -5.20, ML ± 0.70, DV 2.50; PCG, AP -5.50, ML ± 0.40, DV 3.1; LHA rostral part, AP -1.06, ML ± 1.0, DV 4.95; LHA caudal part, AP -2.00, ML ± 1.0, DV 4.95.

To implant an optical fiber in the target brain region for optical stimulation or fiber photometry recording, we drilled 3 to 4 small holes into the surface of the skull and inserted skull screws. Simultaneously, we polished the surface of skull with a cranial drill to increase friction. After drilling the hole at the coordinates corresponding to the target brain region, the optical fiber was slowly lowered into the brain using the stereotactic arm at a speed of 1mm/min. Once the optical fiber was in place, the gap in the skull was first filled with a biocompatible adhesive, and this was followed by primary fixation with light-curing adhesive, and finally the optical fiber was secured to the skull with dental cement.

To retrogradely target suparspinal neurons, we injected viruses into the lumbar segment of spinal cord. For the stereotactic injection into the spinal cord, a dorsal incision was made along the spine at the level of the lumbar segment. The overlying muscles were stripped to expose the lumbar spine, and a laminectomy was performed to expose the spinal cord. The virus was injected into the lumbar spinal cord at a rate of 60 nL/min. After the injection, the micropipette was held in place for a few minutes before it was slowly withdrawn.

**Single-neuron projectome: data acquisition and analysis**

**fMOST imaging and preprocessing**

As outlined in previous descriptions, the fMOST imaging process was carried out. Initially, the brains that had been dissected were subjected to post-fixation in 4% paraformaldehyde (PFA) before being embedded in Lowicryl HM20 resin (Electron Microscopy Sciences, 14340). Subsequently, the brains were imaged in a propidium iodide (PI) filled water bath using an fMOST microscope, achieving a voxel resolution of 0.32 μm × 0.32 μm × 1 μm. The surface of the sample in a coronal plane was then imaged to trace the neurons (GFP) and register the brain (PI channels), after which it was sectioned in 1 μm increments using a fixed diamond knife. This procedure, involving cycles of sectioning and imaging, was repeated until comprehensive imaging of the entire brain sample was accomplished.

We used a proprietary software tool, Fast Neurite Trace (FNT), to specifically trace long-range axon projections in the datasets produced by light microscopy. Initially, the software's "slice2cube" function divides the original image data into more manageable 3D data cubes. At any given moment, approximately eight adjacent cubes around a specified position are automatically loaded into the computer's memory and rendered in 3D for tracing. The tracing process consists of identifying a potential path, scrutinizing it, and expanding the existing neurite tree. User confirmation is required at each step to ensure the accuracy of the tracing process. This method allows for the comprehensive tracing of an axon projection. To ensure quality control, different neurons within the same sample were traced by different team members concurrently, with results cross-validated upon completion. Moreover, each traced neuron underwent an independent review by a separate individual to confirm accuracy. Additionally, we conducted random post-tracing quality checks on each brain sample.

The entire brain, complete with the information of all reconstructed neurons, was registered into the standard Allen CCFv3 using a method previously described. To summarize, we segmented several brain regions as landmarks, using cytoarchitecture references. We then executed a diffeomorphic transformation and symmetric image normalization using Advanced Normalization Tools (ANTs). This process enabled us to acquire transformation parameters based on these landmarks. Subsequently, these transformation parameters were applied to all the traced neurons within the brain sample. This allowed us to remap the reconstructed neuron onto the Allen brain template.

**Neuron exclusion**

We carried out manual checks on all neurons to confirm the accuracy of the tracing. Neurons that been found to have incorrect tracing data and with soma locations outside the lateral hypothalamus were excluded from subsequent analysis, after which a total of 916 neurons located in the LHA was incorporated in our analysis. All the neurons were mirrored to the same hemisphere prior to analysis.

**Parameter calculation and neuron visualization**

Utilizing the standard Allen CCFv3 annotation file as a reference, we computed the soma location, axon length, and terminal number for each neuron in every brain region. This was achieved using a Python library we developed, which is available at <https://pypi.org/project/pyswcloader/>. All neuron visualizations were generated using a self-developed Python package, neuron-vis (<https://gitee.com/bigduduwxf/neuron-vis>).

**Hierarchical clustering of LHA neurons based on axon morphology**

The projections and morphological information of LHA reconstructed neurons can be downloaded from our publicly accessible website (<https://mouse.digital-brain.cn/projectome>).

To obtain the similarity matrix of axon morphology, first we extracted the coordinates of each point from the original morphological file of each neuron, which were referred to by neuron P and Q, respectively. For each point i in the neuron P, we calculated the shortest distance to neuron Q and denoted this value as d(Pi→Q). The same calculation was performed on each point j of neuron Q to obtain the shortest distance to the neuron P, which is denoted by d(Qj→P). Then we calculated a parameter α, the proportion of data points smaller than the mean value in each set of shortest distance, to get the weighted average value according to the following formula:

d(P→Q) = α*mean(d(Pi→Q)) + (1-α)*max(d(Pi→Q))

d(Q→P) = α*mean(d(Qj→P)) + (1-α)*max(d(Qj→P))

The similarity score between neuron P and Q was calculated as the mean value of d(P→Q) and d(Q→P). Then hierarchical clustering using Ward’s linkage was performed on the similarity score matrix to identify the subtypes.

**DTR-mediated cell ablation**

For local inhibition of the LHA, we injected Cre-dependent AAVs carrying diphtheria toxin receptor (DTR) into LHA of the *Vglut2*-Cre mice. For intersectional inhibition of PnO or MSDB projecting LHA neurons, we first injected AAV2-retro-Cre into the PnO or MSDB and then injected AAV-DIO-DTR-mCherry into the LHA of wild type mice. Three weeks after DTR injection, diphtheria toxin (DT, 50 μg/kg, Sigma) was intraperitoneally injected for three consecutive days to deplete the neurons. All the behavioral tests were conducted at least seven days after the DT injection. To confirm the effects of neuronal lesions, mice were sacrificed and perfused.

**Optical-fiber-based Ca2+ recording in freely behaving mice**

After 3 weeks of surgery, fiber photometry recording was carried out by using a commercial device (RWD life science, Shenzhen, China) as previously described. In brief, laser beams of 470 nm and 410 nm were initially launched into the fluorescence cube, then launched into the optical fibers. The 410 nm laser was used for motion control. The emission fluorescence from both the GCaMP and control was collected by a camera at the frequency of 20 Hz. The in vivo recordings were carried out within a custom-made open field box (40 × 40 × 40 cm), with food placed in a corner.

We calculated the value of the photometry signal, denoted as F, using the ratio F_470_/F_410._ We then computed ΔF/F as (F – F_0_) / F_0_, where F_0_ is the median of the photometry signal. Only behaviors with calcium signals exceeding 3 standard deviations (SD) were treated as events. The average of 10 peak ΔF/F and the number of events per minute for each mouse were analyzed.

**Single unit recording and analysis**

The 32-channel drivable electrodes were hand-assembled as previously described (Lin et al., 2006). Before the surgery, the impedance of the tetrodes was adjusted to 300-600kOhm. The tips of the tetrodes were implanted above the LHA and the electrodes were then fixed to the skull with dental cement. Recordings began five days after surgery, during which mice were allowed to move freely in a linear runway (1.5m*0.1m*0.15m). After each recording, the electrodes were advanced by 100-200 um. All signals were acquired using TDT system (Tucker-Davis Technologies, USA), digitalized and sampled at 24 kHz, then bandpass filtered online (0.7-3 kHz). The stored data was converted and then processed using Offline Sorter for single-cell isolation (Plexon, Inc., TX, USA). The corresponding speed was obtained from synchronized videos recorded by the TDT system, which were analyzed using EthoVision XT (Noldus). The correlation between firing rate and speed was analyzed in a manner similar to previous work (Roseberry et al., 2016).

**EMG Recording**

For electromyography (EMG) recording, we performed implantation of customized bipolar electrodes into selected hindlimb muscles to record EMG activity. Electrodes (793200, A-M Systems) were guided by 30-gauge needles and inserted into the mid-belly of the medial gastrocnemius (GS) and tibialis anterior (TA) muscles of the hindlimb. A common ground wire was inserted subcutaneously in the neck-shoulder area. Wires were routed subcutaneously through the back to a small percutaneous connector, which was securely cemented to the skull of the mouse. EMG signals were acquired using a TDT system (RA32 pre-amplifier + RZ2 base processor) controlled by Synapse software (TDT) with a filtration range of 10-1000 Hz. The acquired signals were analyzed using custom-written MATLAB scripts.

**Behavioral experiments and kinematic analysis**

Motor functions were assessed weekly with a locomotor open field rating scale, the Basso Mouse Scale (BMS) (Basso et al., 2006). All BMS behavioral tests were conducted by investigators who were blinded to the treatment groups.

Behavioral tracking of mouse movements was performed offline utilizing DeepLabCut (<https://github.com/DeepLabCut/>). Behavioral tests included overground locomotion of mice either prior to or throughout the recovery period following SCI on an elevated runway. More than 15% of the frames were selected from the behavioral videos of mice in different states to serve as the initial training datasets. To quantify the gait characteristics of mice before and after SCI, frames in the training datasets were manually labeled for the iliac crest, hip, knee, ankle and toe. A ResNet-50 network was trained for 1×10^5^ iterations on the initial training datasets, and the network with the lowest loss value was selected for subsequent analysis through evaluation. Videos representative of different mice were analyzed using the trained ResNet-50 network to obtain the labeled videos and the coordinates of the labeled points.

Kinematic variables were computed using custom scripts in MATLAB (v.R2018a, MathWorks), based on the x, y coordinates generated by DeepLabCut. The whole limb oscillation was evaluated by calculating the angle of the virtual limb linking hip to toe. The speed of toe was calculated by taking one frame from every six frames to determine whether the mouse was in the stance or swing phase, with the threshold being adjusted for each individual mouse. To visualize the movement of mice, we extracted a 3-second segment from the videos of different mice and displayed the labeled points and skeleton lines of their lower limbs.

**Open-field (OF) test Food seeking test and elevated plus maze (EPM)**

The test apparatus consisted of a rectangular box (40 × 40 × 30 cm). A dim light was applied above the field. Before the experiment, the mouse was put into the test room for a 30-minute acclimation period. Subsequently, the mouse was gently placed into the center of the rectangular box for free exploration. The time spent in the center zone (20 cm × 20 cm), as well as the total distance and center time traveled in the whole open field arena were measured over 20 minutes.

Specifically, for the group assigned to the task of food seeking, mice underwent a 24-hour food restriction period prior to the test to enhance their motivation to food seeking during the assessment. During the fasting and testing period, they were maintained at more than 90% of their free-feeding body weight. Food pellets were placed in the center area and could be obtained by a single mouse in the OF for 20 min.

The EPM (Elevated Plus-Maze) apparatus used in our study consisted of a central region (5 × 5 cm), two open-arms (30 × 5 cm), and two enclosed arms (30 × 5 × 15 cm). The apparatus, arranged in a “+” configuration, was situated 50 cm above the floor. At the beginning of the EPM test, the mice were placed in the central area, oriented towards an enclosed arm, and allowed to explore for a duration of 10 minutes.

In addition, we utilized EthoVision XT (Noldus) and custom-written MATLAB scripts to extract the total time that the animals spent in each zone of the open field or each arm of the EPM. Example trajectories were generated using EthoVision XT (Noldus). For both the open field and EPM test, we calculated the velocity by dividing the total distance by the moving time, data for which was also extracted using EthoVision XT.

**SCI Surgical procedures**

The procedure for the T7 and T10 double lateral hemisection and T10 lateral hemisection was performed by a surgical technician. For the T7 and T10 double lateral hemisection, a T7-T10 laminectomy was made over the thoracic vertebrae. We utilized a scalpel and micro-scissors to disrupt the bilateral dorsal columns to make the T7 right-side over-hemisection, to make sure no sparing of the ventral pathways on the contralateral side. We carefully sectioned only the left side of the spinal cord up to the midline at T10. Subsequently, the muscle layers were sutured, and the skin was secured using wound clips. For the T10 lateral hemisection, a midline incision was made over the thoracic vertebrae followed by a T10 laminectomy. The unilateral hemisection was then performed carefully using both scalpel and micro-scissors, avoiding, to the greatest extent, the damage of the spinal cord dura.

**Post-surgical treatments and care of the animals**

Following SCI in adult mice, we administered a subcutaneous injection of 1 mL saline, positioned food on the floor of their cages, and ensured that they were kept warm until they regained consciousness. We undertook manual bladder emptying procedures twice a day and monitored for signs of dehydration and any clinical indicators of pain or infection. In case of a urinary infection, we initiated a course of antibiotherapy (Baytril, 10mg/kg, for 5 days). Any mice exhibiting a body weight reduction exceeding 15% were humanly euthanized.

**Deep brain stimulation**

The deep brain stimulation (DBS) apparatus is composed of bipolar parallel tungsten wires, each with a diameter of 0.1 mm and a length of 5.5 mm. The insulation was removed at a length of 200 μm from the tip. The two electrodes were spaced 2.2 mm apart to bilaterally target the LHA. The DBS electrodes were secured to the head with dental cement. The mice were allowed to recover in their home cages for at least 3 days before starting the behavioral tests.

In the electrical stimulation mapping experiments, a stimulating electrode was lowered into the LHA under ketamine anesthesia. The electric current of -75 μA, 0.5 μs was delivered at a frequency of 45 Hz for 5 seconds, with intervals of 30-60 seconds between trials. The peak-to-peak amplitude and latency of the evoked responses were analyzed from EMG recordings of the right tibialis anterior (TA) muscle. Evaluations were conducted both with and without stimulation. The selected stimulation sites were strategically positioned along the AP axis of LHA at the coordinates of -0.54, -1.04, -1.54, -2.04, and -2.54 mm.

**Close-looped DBS**

Two electrodes for recording and LHA stimulation were implanted under aseptic conditions and general anesthesia. To record the activities of M1 neurons, a 32-channel microelectrode array (4 × 8 array of tungsten wires, spaced 100µm apart) was inserted into layer Ⅴ of the hindlimb area of the primary motor cortex (M1). In addition, a deep brain stimulation (DBS) apparatus was inserted in the LHA. The ground and reference wires from the array were attached to screws fixed to the skull.

During online testing, neural signals from M1 were monitored in real-time. Whenever the value of y=wn exceeded a detection threshold within the range of 0.7 to 0.8, the real-time processor administered LHA-DBS. Conversely, the DBS was halted when it fell to the threshold within the range of 0.2 to 0.3 (Bonizzato et al., 2021).

**Immunohistochemistry and Imaging**

Mice were perfused with phosphate-buffered saline (PBS) followed by 4% paraformaldenhyde (PFA) in PBS. The brain and spinal cord were removed and postfixed in 4% PFA at 4 ℃ for 24h. The tissue was dehydrated in 30% sucrose solution at 4℃ for 24h and sectioned coronally into 40 μm slices with a Cryostat microtome (Thermo Scientific HM525 NX). The sections were washed with PBS (6*10min) and then blocked with a blocking solution (PBS containing 10% donkey serum and 4% Triton X-100) at room temperature for 2h. Then the sections were incubated with the primary antibodies at 4℃ for 24h. Upon completion of the primary antibody incubation, the sections were washed with PBS (6*10min) and then incubated with secondary antibodies at room temperature for 2h. After the 2-hour incubation, the sections were thoroughly washed with PBS (6*10min) and mounted with anti-fade mounting medium with DAPI for imaging. Sections were photographed using the Olympus VS120 virtual slide system.

**QUANTIFICATION AND STATISTICAL ANALYSIS**

All analyses were conducted blindly with respect to experimental conditions, using GraphPad Prism 10.0 software or MATLAB (v2018a). A two-tailed Student’s t-test was employed for single comparisons between two groups. For all other data, one-way or two-way ANOVA was utilized based on the appropriate experimental design. Post hoc analyses were performed exclusively when a main effect reached statistical significance, with P values for multiple comparisons adjusted using Bonferroni’s correction. All data are presented as the mean ± SD unless otherwise stated. ns, not significant; P > 0.05; *P < 0.05; **P < 0.01; ***P < 0.001; ****P < 0.0001. Mice were randomized by litter, body weight, and sex before assignment to treatment groups; no additional specific randomization was implemented for the animal studies.
